# A temporal map of gene expression pattern during zebrafish liver regeneration

**DOI:** 10.1101/677781

**Authors:** Urmila Jagtap, Ambily Sivadas, Sandeep Basu, Ankit Verma, Sridhar Sivasubbu, Vinod Scaria, Chetana Sachidanandan

## Abstract

**Background & Aims:** Zebrafish is increasingly being used to study liver injury and regeneration. However, very little is known about molecular players that respond to injury and participate in liver regeneration. Here we aim to generate a temporal map of gene expression changes at injury and during regeneration of the adult zebrafish liver.

**Methods:** We use a metronidazole-nitroreductase (MTZ-nfsb) based system to selectively ablate hepatocytes in adult zebrafish to create a model for liver injury and regeneration. Through RNA sequencing of liver samples at multiple time points we generate a comprehensive temporal map of gene expression changes during injury and regeneration.

**Results:** Gene expression reveals that soon after injury the immediate early transcription factor MYC induces a battery of genes that respond to the metronidazole-induced ROS by activating oxido-reductase pathways and apoptosis machinery. Upon injury, liver cells down regulate genes encoding complement proteins, bile acid and lipid biosynthesis pathway in a concerted manner. Midway through regeneration, we discover a spike of cholesterol biosynthesis and protein folding machinery genes suggesting an important role for these pathways in liver regeneration.

**Conclusions:** The temporal transcriptomic map of liver regeneration would serve as a framework for further studies in understanding, and for screening for compounds that augment liver regeneration.

**General significance:** Using a hepatocyte specific ablation of zebrafish liver, we create a model of adult liver regeneration. This model was used to generate a comprehensive transcriptomic map of gene expression trends during liver regeneration. This temporal map lays the groundwork to study important events in liver regeneration.

**Highlights:**

- Zebrafish is a valuable model for developing therapeutic strategies to augment liver regeneration
- Liver regeneration in zebrafish is not well studied and pathways poorly understood
- We develop a hepatocyte ablation model of liver injury and regeneration in adult zebrafish
- We generate a comprehensive transcriptomic map of various stages of liver injury and regeneration
- We discover a novel regulation of cholesterol biosynthesis pathways during liver regeneration

**Graphical abstract:** 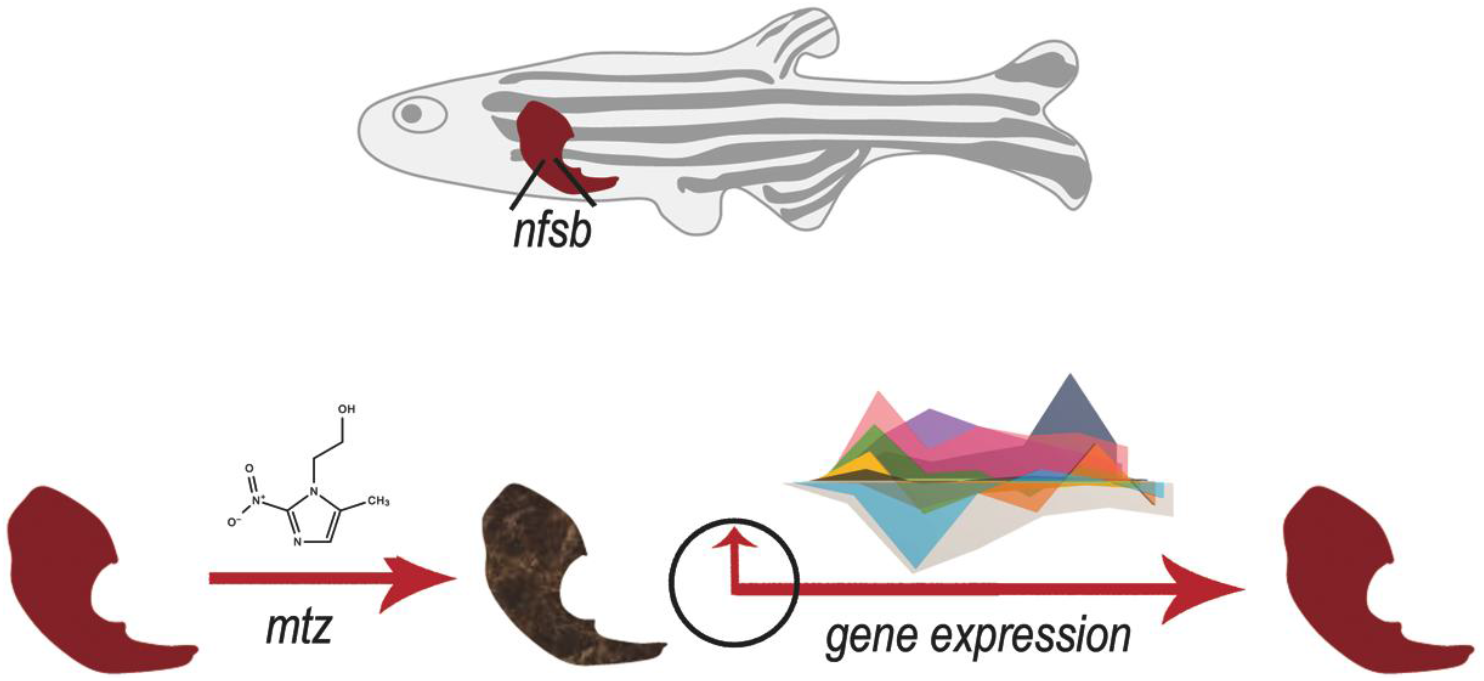

## Introduction

Liver is a highly regenerative organ. However, the capacity and rate of repair is often not sufficient to overcome the loss of cells during liver damage (1–3). Failure to appropriately compensate for the loss in the functional units of liver often leads to multiple organ failure and severe morbidity and mortality (4). Studies on animal models of liver injury are crucial to identify strategies to augment liver regeneration. Most of our knowledge about liver regeneration comes from studies on the partial hepatectomy (PHx) model in mice and rat (5), where 50-70% of the liver is resected. Studies on hepatotoxic drug induced liver injury (DILI) and regeneration in animal models have also led to new insights in the field of liver regeneration (6).

Studies on these models established that the gene expression response of the liver to injury can be categorised into two phases: the ‘immediate early’ and the ‘delayed activation’ phases (7). A number of secreted signalling molecules, growth factors, transcription factors and miRNAs have been shown to have very important role in the regulation of the regenerative response (8). Larger transcriptomic studies on liver injury models have generated datasets of gene expression for comparative analysis as well as identified novel mechanisms (7). A recent study analysed the proteome, epigenome and transcriptome of damaged liver in mice and identified FoxM1 as a key regulator of the regenerative response (9). Recent transcriptomic and metabolomic studies on a mouse PHx model revealed that cell division post-injury causes metabolic remodelling of the liver cells and that hepatocytes adapt their metabolism to changing conditions during regeneration (10). A comparative transcriptomic study of lethal and sub-lethal doses of acetaminophen driven drug induced liver injury (DILI) in mice identified factors that may give adaptive advantage in cases where the liver is able to regenerate and survive the damage (11). In spite of the ability of the liver to recover from loss of hepatocytes by regeneration, in liver diseases the recovery is usually suboptimal and not sufficient. Thus, models of liver regeneration, which can be subjected to drug discovery screens, are needed to discover strategies for therapeutic augmentation of liver regeneration.

Zebrafish has been used extensively to model human diseases and discover small molecules that can ameliorate the disease phenotypes (12). Zebrafish has been used to create liver injury models; in fact, one of the few compounds to reach clinical trials for liver regeneration, PGE2, was discovered from an acetaminophen induced liver injury model in zebrafish (13). Zebrafish studies on the MTZ-nfsb hepatocyte ablation system have revealed an important role for trans-differentiating biliary epithelial cells in supplementing hepatocyte loss, at least in zebrafish (14). Zebrafish liver regeneration studies using the MTZ-nfsb injury model have revealed a number of novel insights into the molecular players involved in liver regeneration (15–17). However, in spite of the growing number of studies on liver injury by hepatotoxic molecules and on regeneration in the zebrafish there are very few transcriptomic maps of the liver injury and regeneration models so far (18, 19). The only such study used a PHx model in adult liver and mapped the liver transcriptome of injured liver upto 24 hours post hepatectomy (20).

Here, we report the first comprehensive chronological transcriptomic map of zebrafish adult liver regeneration using the MTZ-nfsb based hepatocyte ablation system. We developed a protocol for liver injury in the adult zebrafish using low doses and short exposure to MTZ. Total RNA sequencing was performed of the whole liver collected immediately after exposure to MTZ, 12 hours, 24 hours, 4 days, 6 days and 8 days after injury and these were compared to the uninjured liver. We observe that the most dramatic changes in gene expression happen in the hours immediately following damage and demonstrate that key signatures of liver damage, as shown in rodent and fish models previously are present in our data set. We further show that, barring a small subset of genes with dynamic changes in later hours, regeneration is characterized by a slow recovery of the pre-damage state. This data set will serve as the reference point for further studies on the molecular mechanisms of liver regeneration as well as for chemical screens for augmenters of liver regeneration in zebrafish.

## Materials and Methods

### Maintenance of adult zebrafish

Zebrafish (Danio rerio) were bred, raised, and maintained at 28.5°C under standard conditions as described (21). Zebrafish experiments were carried out according to standard procedures approved by the Institutional Animal Ethics Committee (IAEC) of the CSIR-Institute of Genomics and Integrative Biology, India in accordance with the recommendations of the Committee for the Purpose of Control and Supervision of Experiments on Animals (CPCSEA), Govt. of India. The study uses a double transgenic zebrafish line generated by crossing *Tg(fabp10a:GAL4-VP16,myl7:Cerulean) (22)* and *Tg(UAS:nfsb-mcherry)* (23).

### Experimental design

1 year old female zebrafish were exposed to 1.5mM Metronidazole in dark for 3 hours. The fish were washed thrice and kept in fresh water. For RNA sequencing experiments, for each time point, 2 fishes were randomly selected, anaesthetized using tricaine methanosulfonate (0.2%) and liver dissected out using standard surgery protocols for zebrafish.

All other experiments such as histology, ROS estimation, quantitative real time PCR, were performed with minimum 3 adult fishes.

### Other Experiments

All other experimental and analysis methods are described in the Supplementary Material and Methods. Raw sequencing data can be accessed from NCBI Short Read Archive (SRA) - BioProject accession: <u>PRJNA543743</u>.

## Results and Discussion

### A zebrafish model to study liver damage and regeneration

In this study, we use a double transgenic zebrafish line *Tg(fabp10a:GAL4-VP16,myl7:Cerulean)(22)*, *Tg(UAS:nfsb-mcherry) (22, 23)* where bacterial nitroreductase (*nfsb*) is expressed selectively in the hepatocytes. Upon treatment with the pro-drug Metronidazole (MTZ) the *nfsb* enzyme converts it into a cytotoxic product causing accumulation of reactive oxygen species (24) and death of hepatocytes. The MTZ is then removed and the liver is allowed to recover and regenerate (Fig.1A). mCherry fused to *nfsb* allows for live detection of liver loss and regeneration over time. We optimized the concentration of MTZ in 1-year-old adult zebrafish by treating animals with various concentrations of the drug: 1.5mM, 2.5mM and 5mM for 3 hours in water. The fish were transferred to clean water and their survival was monitored over 8 days. 2.5mM and 5mM MTZ treatment for 3 hours was found to be lethal. However fish treated with 1.5mM MTZ survived and appeared to be healthy. Thus, this treatment regimen was selected for all further experiments.

**Figure 1:**
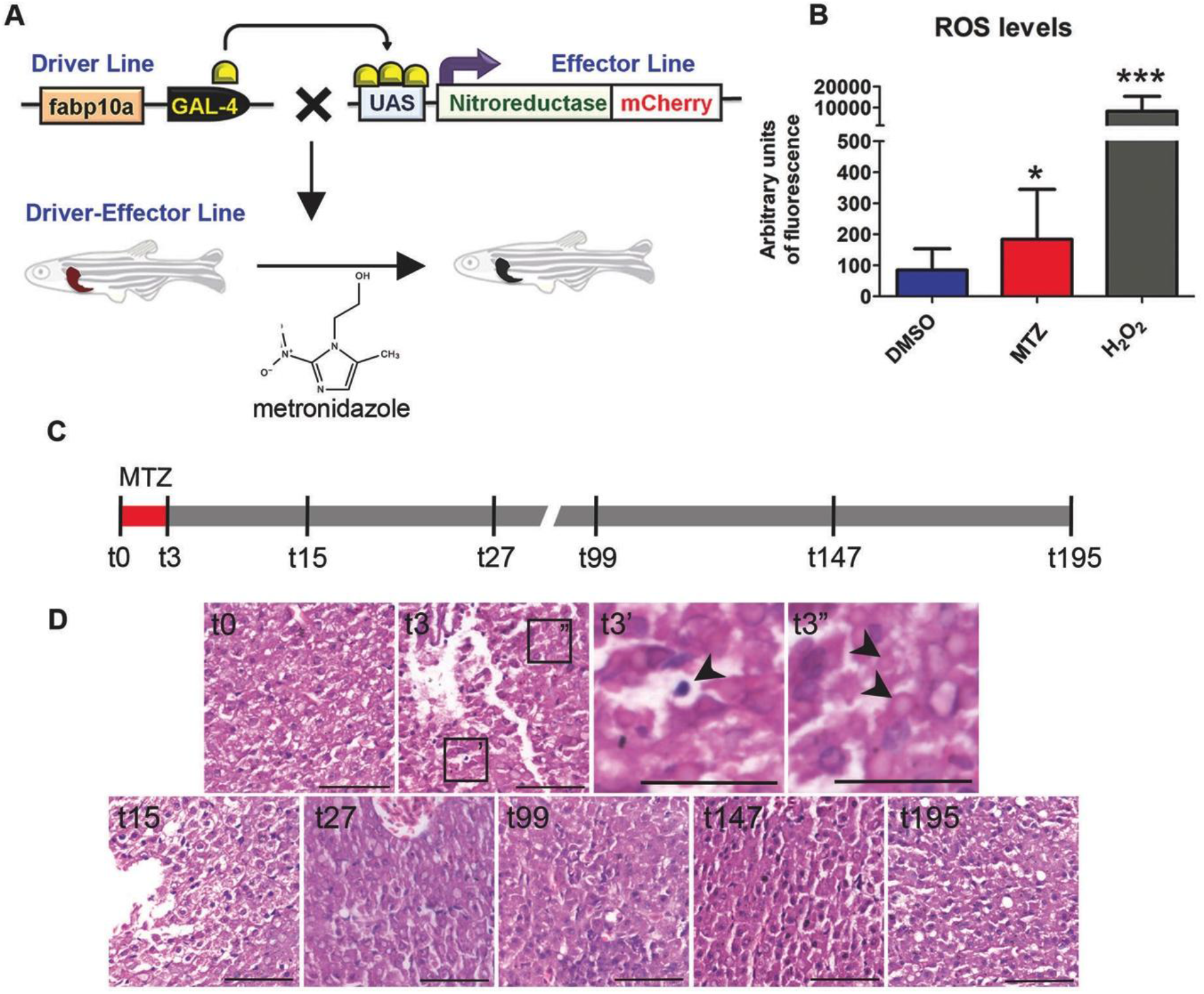
A zebrafish model to study liver damage and regeneration. (A) Schematic depiction of double transgenic zebrafish line with a driver line: *Tg(fabp10a:GAL4-VP16,myl7:Cerulean)* and effector line: *Tg(UAS:nfsb-mcherry)*. The double transgenic line expresses nitroreductase-mCherry specifically in the hepatocytes. Nitroreductase converts the pro-drug Metronidazole into a cytotoxic product, thereby causing hepatocyte specific ablation. (B) DCFDA assay for ROS detection in DMSO, MTZ and H_2_O_2_ treated livers. H_2_O_2_ served as a positive control. (C) Timeline for RNAseq experiments. (D) Histology analysis of each time point of liver injury and regeneration. Black arrowhead marks the presence of pyknotic nuclei (t3’) and enucleated cells (t3”) at t3. There is a a gradual restoration of architecture by t195. Scale bar: 50µm, Magnification: 40X.

We collected livers at various time points following MTZ treatment: immediately (t3), 12 hours (t15), 24 hours (t27), 4 days (t99), 6 days (t147) and 8 days (t195) following MTZ treatment. Untreated (t0) fish served as control (Fig. 1C). We stained sections of the liver with Hematoxylin and Eosin (H&E) and performed a qualitative histological analysis. In zebrafish, the untreated liver is packed with hepatocytes with the occasional portal vein or biliary duct (25). The MTZ treated livers (t3) developed white spaces between the packed parenchyma, indicating edema. There was also a large number of pyknotic nuclei (Fig. 1D t3’ arrowhead) and enucleated cells (Fig. 1D t3” arrowheads) present. These signs of liver injury peaked at t15 and reduced in t27. By day 4 and 6 there was further improvement in the hepatic architecture and an increase in number of nuclei indicating proliferation. By day 8 the liver structure resembled the uninjured liver (t0) with evenly sized and well-packed hepatocytes (Fig. 1D). Since activation of pro-drug MTZ by the bacterial *nfsb* is known to release ROS we quantified and compared the formation of ROS in liver extracts at t0 and t3 using the DCFDA assay. We observed a 3-fold increase in ROS levels in t3 compared to t0 (Fig. 1B). Hydrogen peroxide, the positive control, showed a 10,000 fold increase in ROS levels.

### Differential gene expression in the liver during regeneration

To generate a transcriptomic profile of liver damage and regeneration in zebrafish we collected RNA from livers at time points, t0, t3, t15, t27, t99, t147 and t195, in duplicates and performed total RNA sequencing. The sequencing was carried out using Illumina Hiseq 2500 platform to an average depth of 100 million reads which were then mapped to zebrafish genome reference assembly version 10 (Zv10). 79-90% alignment was achieved using the splice-aware aligner, STAR (Fig. 2A).

**Figure 2:**
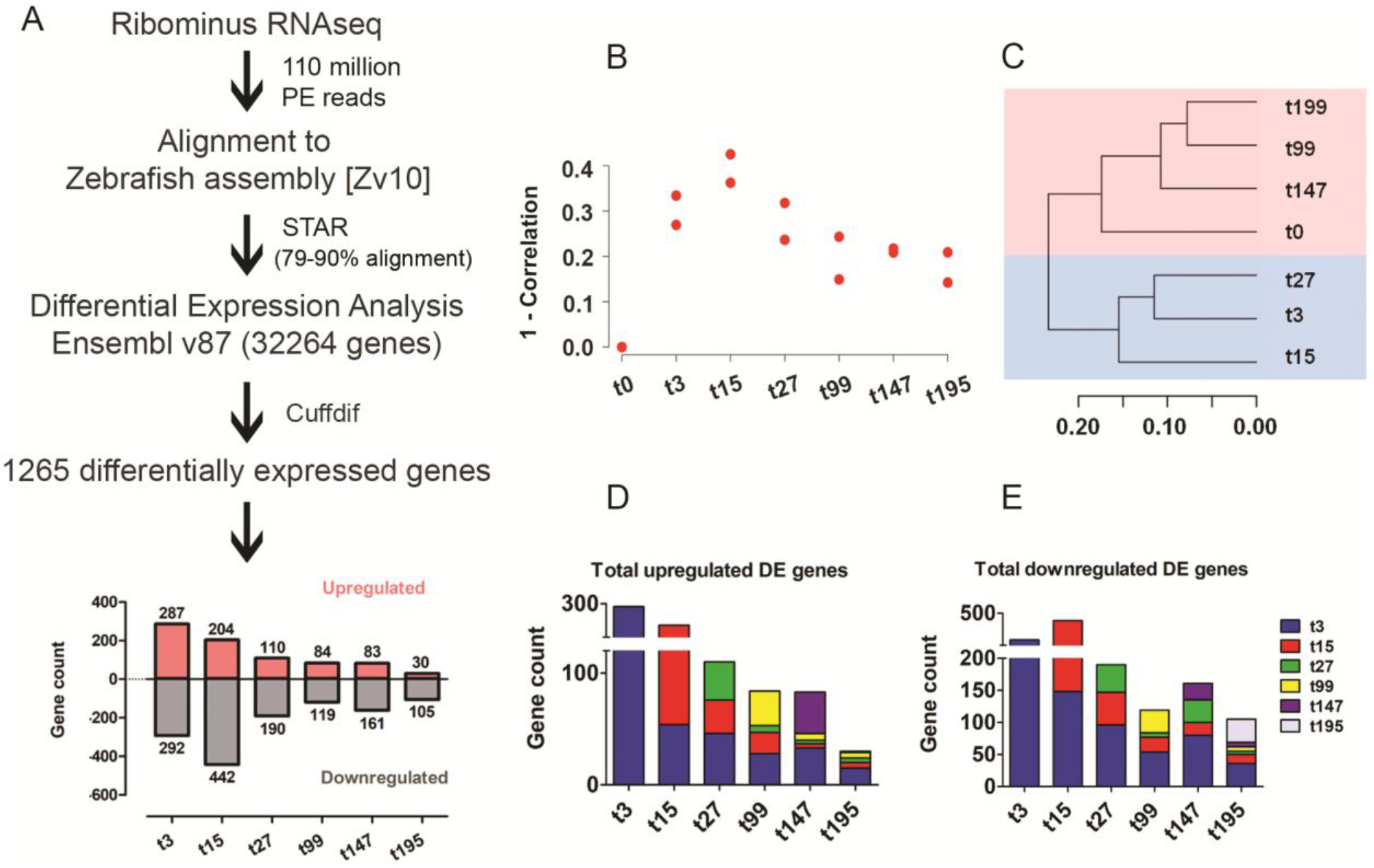
RNAseq analysis workflow. (A) Pipeline used for filtering RNAseq reads. Alignment to ZV10 and Cuffdiff analysis led to 1265 genes with differential expression in at least 1 time point when compared to t0. Orange and grey bars indicate the number of genes up regulated and down regulated respectively at a specific time point. (B) The Pearson correlation plot testing the concordance between each replicate with the normal livers (t0). (C) Hierarchical clustering for all samples using average linkage clustering of a distance matrix obtained from 1-Pearson Correlation as the dissimilarity index. The light red background indicates ‘damage’ phase and the light blue indicates ‘regeneration’ phase. Temporal and differential contribution in terms of (D) induction and (E) repression of DEGs at specific time points.

All samples were compared to t0 using CuffDiff and the differential expression analysis used zebrafish reference transcript annotation model of Ensembl v87. After filtering for genes that had an FPKM>1 and were differentially expressed (p value < 0.05) in at least one time point compared to t0 (Table S1) we identified1265 protein coding genes (Fig. 2A). All further analysis was performed with this gene set.

To assess the global transcriptomic concordance between samples, Pearson correlation was computed between the gene expression profile (N=1265) of each time point (the duplicates were analysed separately) and an average of t0 (Fig. 2B). The t3 liver showed a significant deviation from t0, however, t15 had the minimum r value compared to t0, indicating minimum similarity with normal liver. This suggested that the effect of MTZ persisted till t15. The concordance with t0 improved from t27 onwards to reach a minimum deviation at t195 suggesting a gradual restoration of the healthy liver transcriptome with time (Fig. 2B). We observed very little divergence between the duplicate samples of each stage suggesting low variability between them (Fig. 2B). We performed hierarchical clustering of each sample using average linkage clustering of a distance matrix obtained from 1-Pearson Correlation as the dissimilarity index and obtained two major clusters: one containing t3, t15 and t27 and the other containing t0, t99, t147 and t195 (Fig. 2C). This further demonstrated two phases we classify as ‘damage’ and ‘regeneration’. Clustering of the later phase with t0 indicated recovery of ‘normal’ liver gene expression and in turn, the functions.

We divided the 1265 differentially expressed genes (DEGs) into up regulated and down regulated classes within each stage, in comparison to t0. We observed that the maximum number of genes induced or repressed were at t3 indicating response to liver injury (Fig. 2D, blue block). A further set of genes was differentially regulated at t15 (Fig. 2D, red block) reinforcing that response to injury lasts at least 12 hours post removal of the pro-drug (Fig. 2D&2E). A smaller set of additional genes were induced or repressed at t27, t99 and t147. The minimum differential expression with respect to t0 was observed in t199, again suggesting a restoration of homeostasis (Fig. 2D&2E).

### Patterns of gene expression changes in the liver during damage and regeneration

The fold change for each of the 1265 DEGs with respect to t0 was visualized as a heat map (Fig. 3A). Applying weighted gene correlation network analysis (WGCNA) to cluster genes with closely related expression profiles revealed 8 different discernible modules of gene expression (Fig. 3A). We traced the expression dynamics of each of the 8 modules. Module 1 and module 5 consist of genes that are rapidly down regulated during treatment and do not recover within the time studied (Fig. 3C and 3G). Module 2 genes show a sharp increase in t3 followed by gradual recovery to pre-damage conditions (Fig. 3D). Genes of module 3 are down regulated sharply with minima at t15 and increase to peak at day 6 (Fig. 3E). Module 4 consists of genes that spike at t3 and drop sharply at t15 (Fig. 3F). Module 6 includes a very tightly regulated set of genes that are induced upto an average 4 fold at t15 and return to their original levels by t27 (Fig. 3H). Module 7 genes appear to follow a cyclic pattern with sharp drop at t3, gradual increase to peak at t99 followed by gradual decrease to minimum levels at t147 and induction again to t199 (Fig. 3I). The final module, module 8, consists of genes with a broad expression pattern of reduction at t3 and induction at t15 (Fig. 3J). The final group of genes shown in the heatmap (grey bar) are genes with no distinctive pattern to their regulation (Fig. 3A). The genes in each of the modules when further assessed for enrichment of biological pathways using KEGG database revealed enrichment of various metabolic pathways (Fig. 3B).

**Figure 3:**
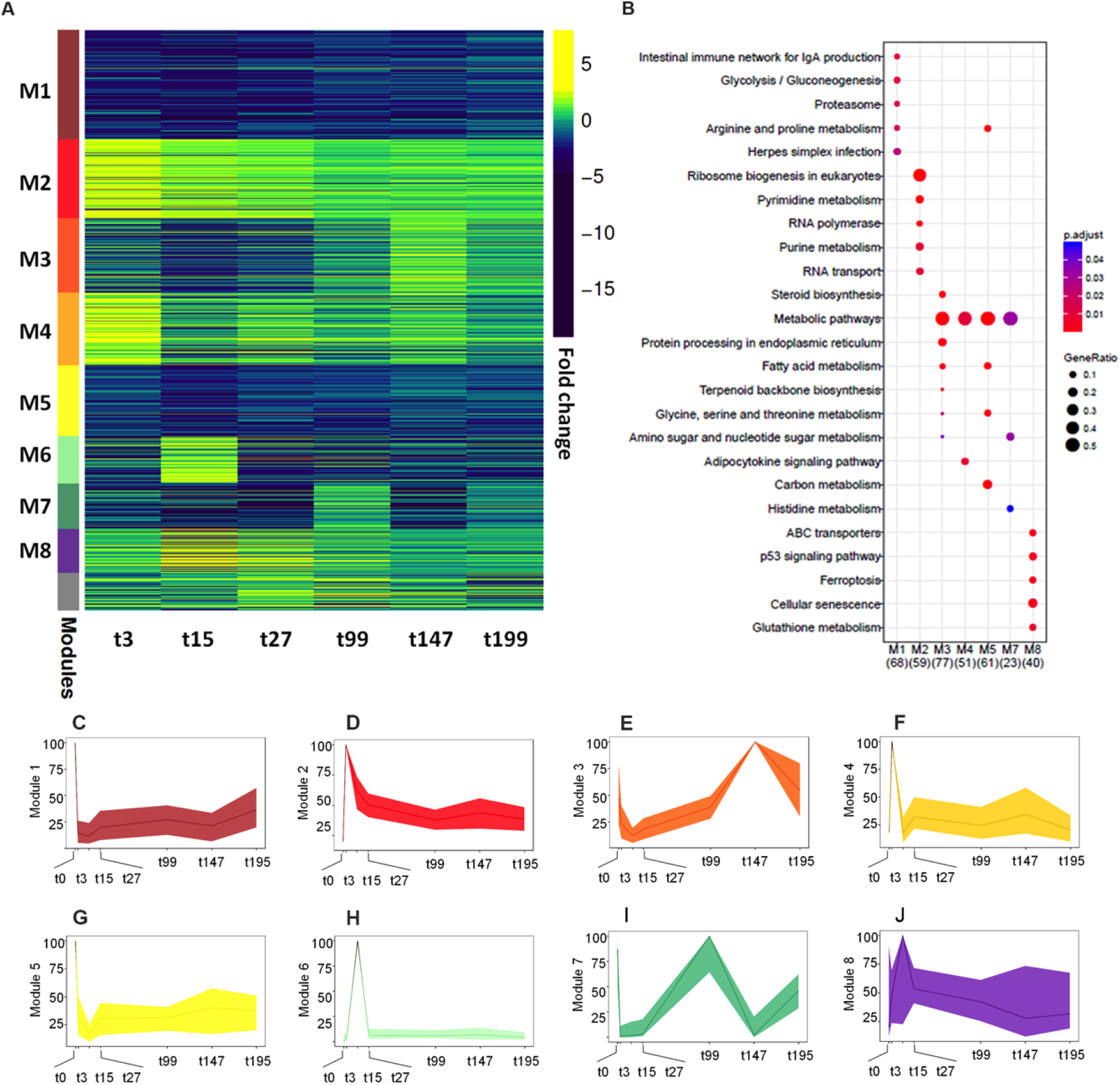
Patterns of gene expression changes in the liver during damage and regeneration. (A) WGCNA analysis categorising the 1265 DEGs into 8 differential modules. The fold change with respect to t0 is plotted. Deep blue indicates up regulation and yellow indicates down regulation. The trace of each module corresponding to its expression pattern (C-J) is plotted across all time points along with the (B) biological theme enrichment for each module.

We utilized the WGCNA measure of intra-modular connectivity (kME) to identify hub genes for each module. Hub genes represent those genes that are highly correlated with the module expression (first principal component (PC1) scores) and therefore potentially regulating the module functions. A total of 126 genes, the top ten percent of genes with highest ranking kME within each module were identified (Table S2). Of these genes, 41 were previously reported in various regenerating tissues in zebrafish e.g. caudal fin, heart and hair (26–28) suggesting a common signature of injury and regeneration. Of the 126 zebrafish genes, 11 had human orthologues. Using the Cytoscape plugin GeneMANIA, a network based on physical, genetic and correlation-based interactions was plotted for the human orthologues (Fig. S1). Clustering of the genes based on their connectivity to each other revealed that the human orthologues of co-expressed zebrafish genes also show high interconnectivity such that genes of the same module (in same colour) cluster together in connectivity groups. This attests to the relevance of zebrafish liver regeneration studies to human liver regeneration.

We broadly categorised the total 1265 DEGs into up and down-regulated categories based on their response at t3 vs t0 and performed a functional annotation of statistically overrepresented classes for these genes at all time points. Based on the number of genes, p value (with Benjamini correction) and fold change at each time point, an alluvial plot of differentially regulated biological processes was plotted using Sankey software (Fig. S2). The height of each block represents the percentage of genes of the pathway in the dataset. Black borders to the block indicates significant p values (<0.05). The KEGG analysis (Fig. 3B) and the alluvial plot (Fig. S2A) highlight the induction of transcriptional and translational machinery immediately after injury. The largest class of genes down regulated upon damage (Fig. 3B and Fig. S2B) belong to metabolic pathways reiterating importance of metabolic regulation in liver damage and regeneration.

### Liver injury and regeneration causes dynamic changes in liver function

Hepatocyte ablation had dramatic effects on liver gene expression, with 7 modules out of 8 cataloguing sharp damage-associated changes. Modules 1, 3, 5 and 7 showed a steep decrease in t3 and t15. We performed gene ontology analysis on this combined group of genes with the DAVID annotation tool and found that the highest enrichment group contained genes of the oxidation-reduction processes. Although, MTZ is known to cause cell ablation through induction of ROS (29) the canonical ROS responsive genes e.g. SOD or catalase are not among our 1265 DEGs. Gene ontology analysis of module 4, genes that spike sharply at t3, identified a group of oxidation-reduction genes (Fig. 4A). Some of the most striking induction (with a 13 to 49 fold) was in genes also identified in an MTZ-*nfsb* based neuronal ablation model in zebrafish (29) e.g.*loxl2a*, *bbox1*, *tdo2* and *cyp2r1*. This non-canonical oxido-reductase response might be a signature of the MTZ-*nfsb* based cell ablation.

**Figure 4:**
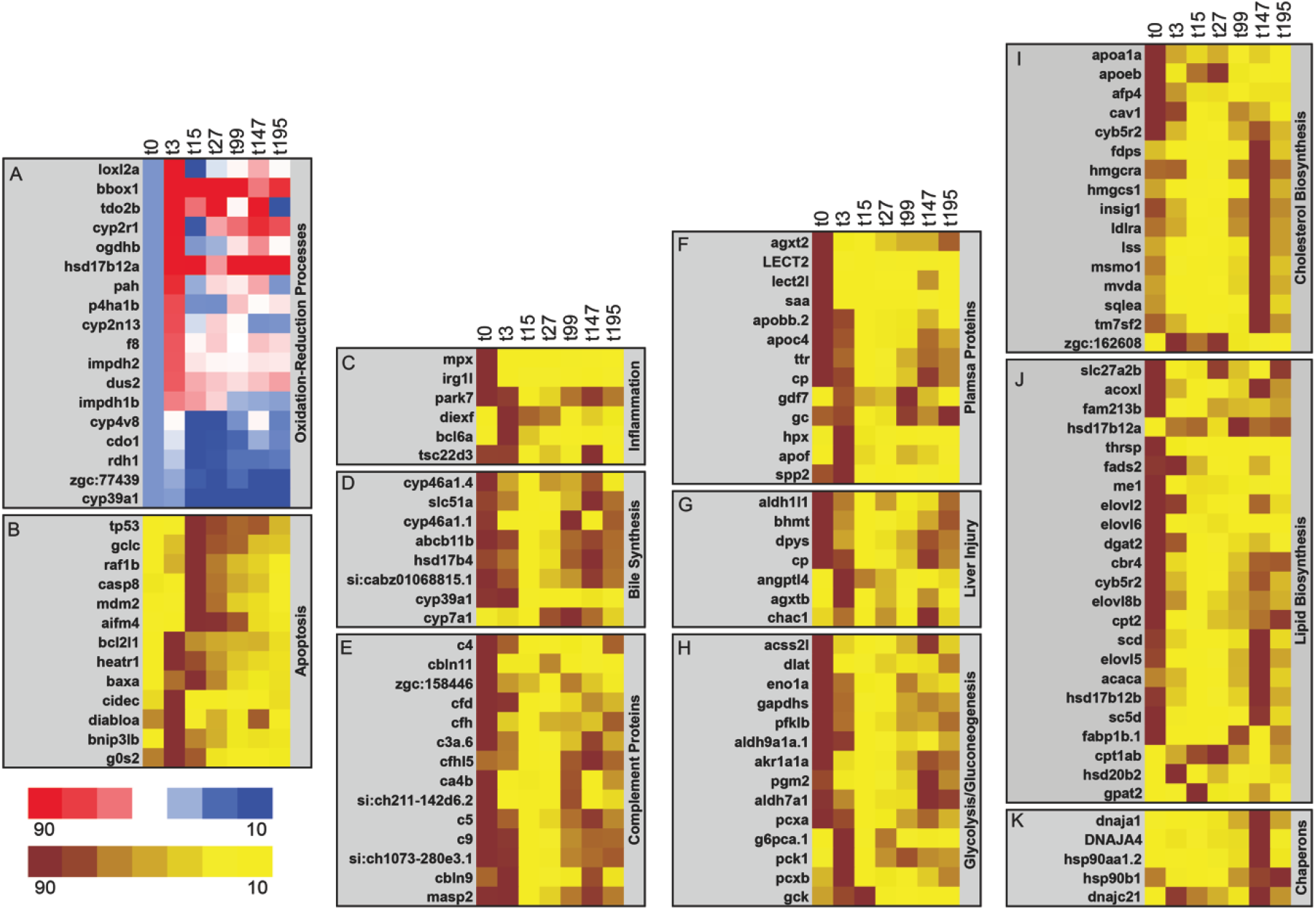
Liver injury and regeneration causes dynamic changes in liver function. (A) Induction of non-canonical oxido-reductase genes as a signature response of the MTZ-*nfsb* based cell ablation. Fold changes in FPKM of each gene at each time point normalized to t0. Red shows up regulation (top 10 percentile) and blue represents down regulation (bottom 10 percentile). (B-L) Average FPKM value of each time point for each gene was used to create the heat map. Maroon indicates the top 10 percentile and yellow indicates the bottom 10 percentile of the values in each row. Patterns of (B) apoptosis markers, (C) inflammation genes, (D) bile synthesis, (E) complement function, (F) liver plasma proteins, (H) glycolysis/gluconeogenesis, (I) cholesterol biosynthesis, (J) lipid biosynthesis and (K) chaperons are plotted.

Drug induced liver injury leads to apoptosis, necrosis or ferroptosis in the liver depending on the type of drug (30); a recent study on the MTZ-*nfsb* model of liver injury in embryonic zebrafish showed that apoptosis dominated (31). We analysed apoptosis genes in our DEGs dataset. At t15 we found a 3-fold induction of p53, a marker of DNA damage and apoptosis likely induced by the MTZ induced DNA damage. Other pro-apoptotic markers such as *baxa*, *caspase8* (*casp8*) and *aifm4* and the p53 regulator mdm2 are also induced by t15 (Fig. 4B). The induction of *p53*, *casp8* and *mdm2* was confirmed by quantitative RT-PCR of RNA from an independent experiment with multiple adult fish livers (Fig. S3A). In keeping with the anti-inflammatory nature of apoptosis, and as shown in the zebrafish embryonic MTZ-nfsb liver injury model (31), we find a down regulation of pro-inflammatory molecules such as the *mpx* (neutrophil marker) while anti-inflammatory molecules such as bcl6a (32) and tsc22d3 (Fig. 4C) are induced upon damage.

Around 431 genes belonging to the combined modules 2, 4 and 6, including apoptosis genes are induced at t15. To identify the transcription factors that regulate this response to liver injury we applied a gene set enrichment analysis tool ENRICHR (33, 34) on these genes. The highest enrichment (18% of the genes) was of the *MYC* and *MAX* (Myc associated factor X) target genes (Fig. S4) (Table S3). The immediate early gene MYC, known to be induced within 30 mins of PHx, has well-established roles in the initiation of cell proliferation (35, 36).

We analysed the effect of liver injury and regeneration on important functions of liver such as bile, complement and other plasma protein synthesis, glucose and lipid metabolism. Previous studies show that hepatocyte ablation in the zebrafish MTZ-*nfsb* model causes a collapse of the biliary network. In keeping with this, we found a concerted reduction in the expression of genes involved in bile synthesis at t3 (6 days) (Fig. 4D). We validated expression patterns of *abcb11b* and *cyp7a1* by quantitative RT-PCR (Fig. S3B, S3C). Synthesis of complement proteins and other plasma proteins is an important function of the liver. We found a consistent down regulation of complement coding genes (Fig. 4E) and a number of other plasma proteins (Fig. 4F) at t3 and t15. Expression of *c3a*.1, *c4* and *c5* were validated by quantitative RT-PCR (Fig. S3D-F). However, we also found strong induction of angptl4 and agxtb (Fig. 4G) as previously reported in zebrafish (20) and human (37).

Glucose and lipid metabolism are important functions of the liver. We found that glycolysis and gluconeogenesis genes were predominantly down regulated at t3 with gradual recovery (Fig. 4H). A gene ontology analysis of genes of modules 1,3,5 and 7 together showed that lipid biosynthesis is a significant subset in this group. The lipid biosynthesis genes were predominantly down regulated at t3 and remained down regulated through the time course (Fig. 4J). Cholesterol biosynthesis genes also showed sharp down regulation upon injury but there was a sharp spike at t147 (Fig. 4I). Many other liver function genes in module 3 e.g. bile synthesis, and complement proteins (Fig. 4D-E) also exhibited this spike at t147. We performed GO analysis using DAVID software on module 3 and found that most enriched class in this module coded for endoplasmic reticulum proteins including chaperones (Fig. 4K) suggesting a translational induction to accompany the transcriptional peak. Compensatory proliferation during liver regeneration is known to cause overgrowth of liver before pruning calibrates the size of liver to the body size (38, 39). The pattern of gene regulation we discovered suggests the existence of a similar functional overcompensation during liver regeneration.

We created a concise map by plotting the average fold change of all genes in a process (Fig. 5). Figure 5 summarizes the intriguing patterns of gene expression dynamics we discovered from this study. These changes during the 8 days of liver injury and regeneration may shed new light on pathways of importance to liver regeneration.

**Figure 5:**
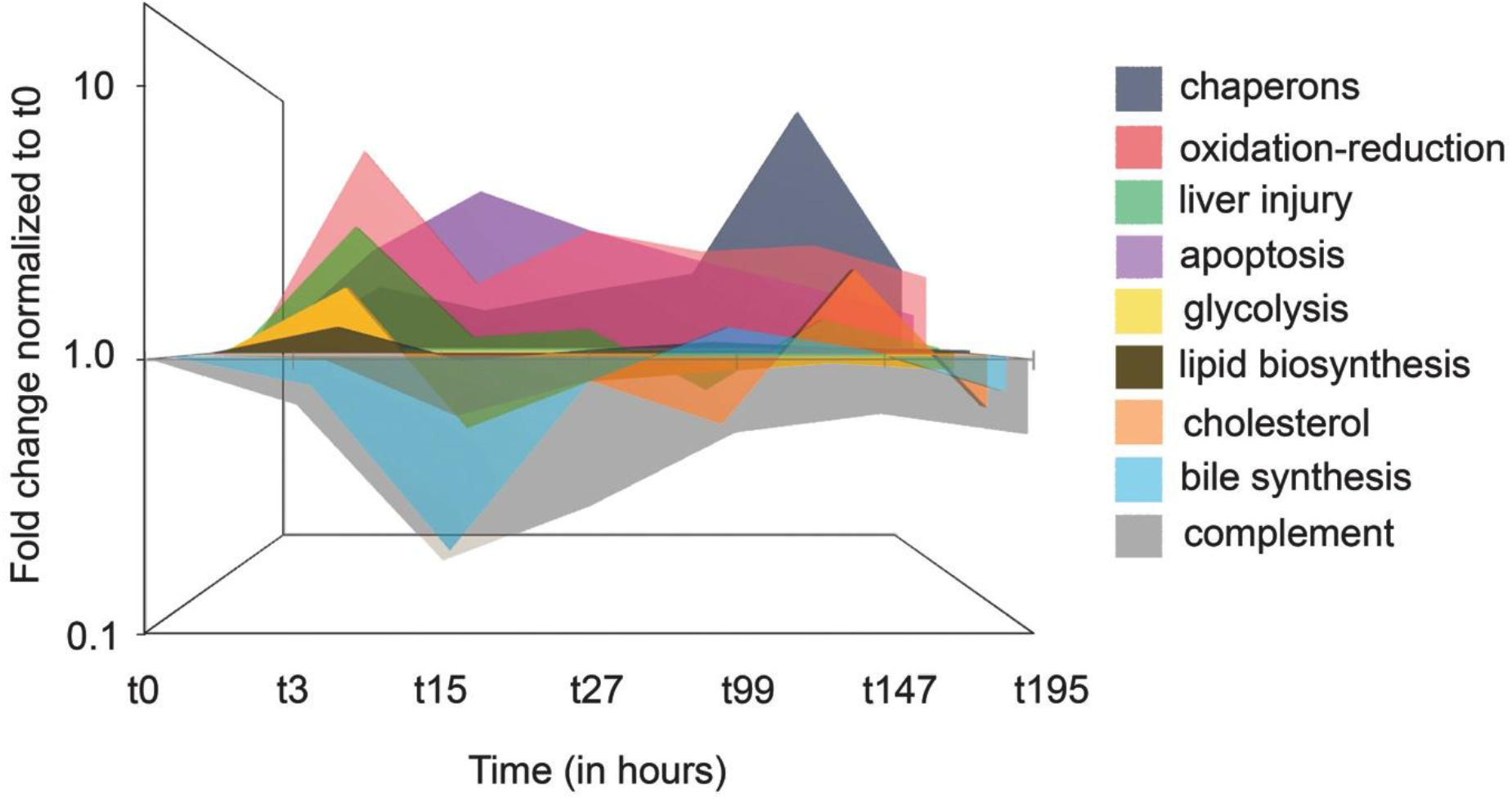
Temporal map of processes differentially regulated during liver injury and regeneration. The fold change for each gene was calculated with respect to t0. An average of fold change of all genes in a specific category was plotted as a function of time.

## Conclusions

Upon injury liver has to rapidly compensate to meet the metabolic demands of the body while repairing the damage to recover full function in the long term. Although a very efficient process, during serious loss of tissue or liver disease, this process falls short. Model organisms such as zebrafish play an important role in understanding and improving the outcome of regeneration. Here we developed an adult model for liver injury and regeneration using the MTZ-*nfsb* system for hepatocyte specific cell ablation. We generated a comprehensive high-resolution transcriptomic map of the gene expression changes after liver injury and until 8 days post damage. The transcriptomic map reveals a surprising down regulation of expression of large number of liver function genes. We discovered that many liver function genes, especially, cholesterol biosynthesis genes transiently spike at 6 days post damage perhaps indicating a functional overcompensation before the liver returns to homeostasis. We speculate that this over-activation of lipid biosynthesis at day 6 has a functional role in achieving successful liver regeneration.

In summary, our study provides a comprehensive system-level analysis of gene expression dynamics during liver regeneration in zebrafish. The temporal map highlights the processes that are regulated tightly and dynamically during liver injury and regeneration suggesting important role in the process of recovery. This transcriptomic temporal map would lay the groundwork for mapping and understanding liver regeneration in zebrafish and could serve as a guiding tool for discovery of small molecules that augment liver regeneration.

## List of abbreviations

MTZ-nfsb: metronidazole-nitroreductase
PHx: partial hepatectomy
DILI: drug induced liver injury
Danio rerio: Zebrafish
DAVID: Database for Annotation Visualisation and Integrated Discovery
kME: module membership scores
MTZ: Metronidazole
H&E: Hematoxylin and Eosin
ROS: reactive oxygen species
DEGs: differentially expressed genes
GO: Gene ontology
MAX: Myc associated factor X
kME: intra-modular connectivity

## Authors’ contributions

**Urmila Jagtap**; Conceptualization, Methodology, Data curation, Formal analysis, Investigation, Validation, Visualization, project administration; **Ambily Sivadas**, Data curation Investigation, Methodology, Software, Formal analysis; **Sandeep Basu**, Methodology, Investigation; **Ankit Verma**, Investigation; **Sridhar Sivasubbu**, Resources; **Vinod Scaria**, Resources, Supervision; **Chetana Sachidanandan**, Conceptualization, Formal analysis, Visualization, Funding acquisition, Resources, Supervision, Writing – original draft, Writing – review & editing*

## Acknowledgements

We thank Kausik Chakraborty, Beena Pillai and Manikandan Subramanian for their critical comments and suggestions during discussions. We thank Tony Jacob and Rajni Yadav (AIIMS, New Delhi, India) for help with analysis and Sarah Iqbal for help with quantification of the histology images. Rijith Jayarajan and KV Shamsudheen helped in library preparation and RNA sequencing runs. We acknowledge all members of the zebrafish facility, especially Shashi Ranjan. Disha Sharma helped in making the data publically available.

