## Supplementary figures and methods for "A temporal map of gene expression pattern during zebrafish liver regeneration"

### MATERIALS AND METHODS

**Tissue histology:**

Liver isolates were fixed in formalin and embedded in paraffin wax. 5-6µm thick sections were cut using MRS3500 (Histo-Line Laboratories). The sections were placed onto the ploy L-Lysine coated slides for Hematoxylin and Eosin staining. Imaging was done at 40X using the Olympus BX61microscope.

**RNA extraction:**

Dissected livers were fixed in RNAiso Plus (Takara, 9109) and total RNA isolation was isolated as per protocol. All RNA samples were treated withDNAse and purity was assessed by 260/280nm absorbance ratios in Nanodrop (ND-1000 UV/vis) and quality was monitored by running samples on 1.5% Agarose electrophoresis. Only samples with 260/280 ratio ~2.0±0.1 were used for library preparation, for RNA sequencing run and for RT-PCR based validation experiments.

**ROS measurement:**

The livers were homogenised, extracts centrifuged at 1000rpm for 10 minutes and the supernatant was incubated with 2',7'–dichlorofluorescindiacetate (H2DCFDA) (Invitrogen-ab113851) for 30 minutes at 37$^{o}$C. Fluorescence was measured at 485/535nm on multimode plate reader Tecan-infinite M200PRO. Background fluorescence was subtracted by measuring fluorescence in identical conditions without the liver extract.

**Real time quantitative PCR:**

Approximately 1 µg of RNA was used for cDNA synthesis using a QuantiTect Reverse Transcription kit (Qiagen, 205313). The amplification factor of each primer set was determined by PCR amplification with serial dilutions of cDNA and only primer pairs with ~100% amplification efficiency were used in the study. Quantitative real time PCR was carried out as described by manufacturer using Sybr green (Roche, 06924204001) on

The *LightCycler* 480 System. Each experiment was performed on minimum three independent adult fishes. Analysis on normalized data used the $2^{-\Delta\Delta}$ CT algorithm (1). All genes were normalized against GAPDH unless mentioned otherwise. p-values were calculated by performing unpaired Student's *t* test. All primers used in this study are listed in Supplementary Table- 4.

**RNA sequencing:**

The RNA-seq library was prepared using 1µg of RNA sample by using a standard protocol (Illumina Incl. USA). High throughput RNA sequencing was performed on HiSeq 2500 platform with a minimum depth of 100 million reads. A paired end strategy (35 cycles) was applied. The data obtained was subjected to Illumina quality control (QC) procedures. The raw paired end sequencing reads of 150 bp length were quality trimmed with a threshold of Q30 followed by adapter removal using Trimmomatic (2). Further, the processed reads were aligned to zebrafish genome reference assembly version 10 (Zv10) using STAR alignment tool (3). Differential expression analysis was performed with Cuffdiff (4) using reference gene annotation from Ensembl v87 with an adjusted p-value i.e. q-value less than 0.05. The fastq files have been uploaded on the SRA data archive with the Project ID PRJNA543743.

**Correlation plot and Hierarchical clustering**

The correlation plot was constructed based on Pearson correlation score of the average gene expression profile of each regeneration time point compared to the untreated t0 stage. The hierarchical clustering analysis of the time points were performed using average linkage clustering of a distance matrix obtained from 1-Pearson Correlation as the dissimilarity index. The above plots were generated using R.

**WGCNA analysis for module development and GO enrichment plot**

The expression data of the differentially expressed genes (n=1265) from 14 samples were utilized for weighted gene co-expression network construction using the WGCNA package. We used a soft thresholding power of 14 for calculating adjacency towards construction of an unsigned weighted correlation network. Using a dynamic tree-cutting algorithm and module merging threshold function at 0.25, we identified eight modules among the differentially expressed genes. The expression pattern for each module was plotted as a function of time. Further, Database for Annotation, Visualisation and Integrated Discovery (DAVID), version 6.8, was used for the KEGG enrichment analysis of each module (5,6). The processes with Benjamini correction <0.05 were considered to be enriched. The gene set enrichment analysis for transcription factor target enrichment was performed using EnrichR (7,8).

**Module enrichment patterns**

The median, first and third quartile expression pattern for each module have been represented as a line plot using R. For this purpose, the individual expression values were normalized to the highest expression value for each gene. The significantly enriched KEGG pathways within each module were visualized in the form of a network using the ClueGOCytoscape plugin (9).

**Sankey plot for up and down regulated genes**

All the DEGs; up and down separately; with respect to t0 at each time point were subjected to gene enrichment analysis using DAVID. Only the enriched terms with a Bonferroni correction <0.05 were visualized as an alluvial plot (10). The pathway node height signifies the percentage of genes enriched in each stage. The nodes with thick boxes indicate a statistically significant enriched pathway at a given time point. The pathway nodes within a time point are arranged in the increasing order of the FDR-corrected p values.

**Hub genes' network**

Intra-modular connectivity (kME) was calculated as the absolute Pearson correlation coefficient between the gene's expression and the module's PC1 scores. Top ten percent of the genes with highest ranking kME within each module were identified as hub genes The human orthologs of these hub genes were utilised to predict comprehensive interaction networks using GeneMANIA Cytoscape plugin using multiple levels of evidence ranging from physical interactions, genetic interactions and correlation-based interactions (11,12).

### SUPPLEMENTARY FIGURES

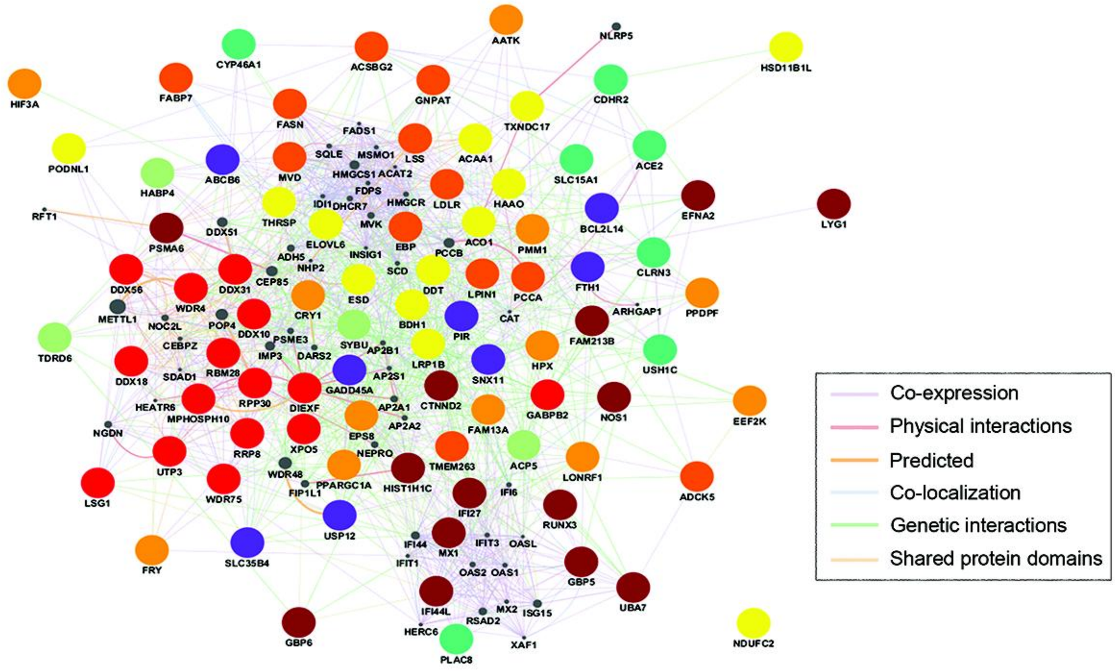


**Figure S1: Hub genes in liver regeneration.** Identification of novel hub genes associated with individual modules using a weighted gene co-expression network to identify hub genes that are highly interconnected with nodes in each module. Top ten percent of the genes with highest intra-modular connectivity (kME) within each module were identified as the hub genes. Using a Cytoscape plugin, Genemania an interaction network of the human orthologues of the zebrafish hub genes, based on their physical, genetic and correlation based interactions were plotted. The colour of each bubble represents the respective module from Figure 3. The connecting lines represented different interactions: purple: Co-expression, light orange: physical interactions, orange: predicted interactions, light blue: co-localization, light green: genetic interactions and light yellow: shared protein domains. The grey circles show 20 genes (not in the our DEGs) related to the hub genes network with their size directly proportional to their connectivity to the input dataset.

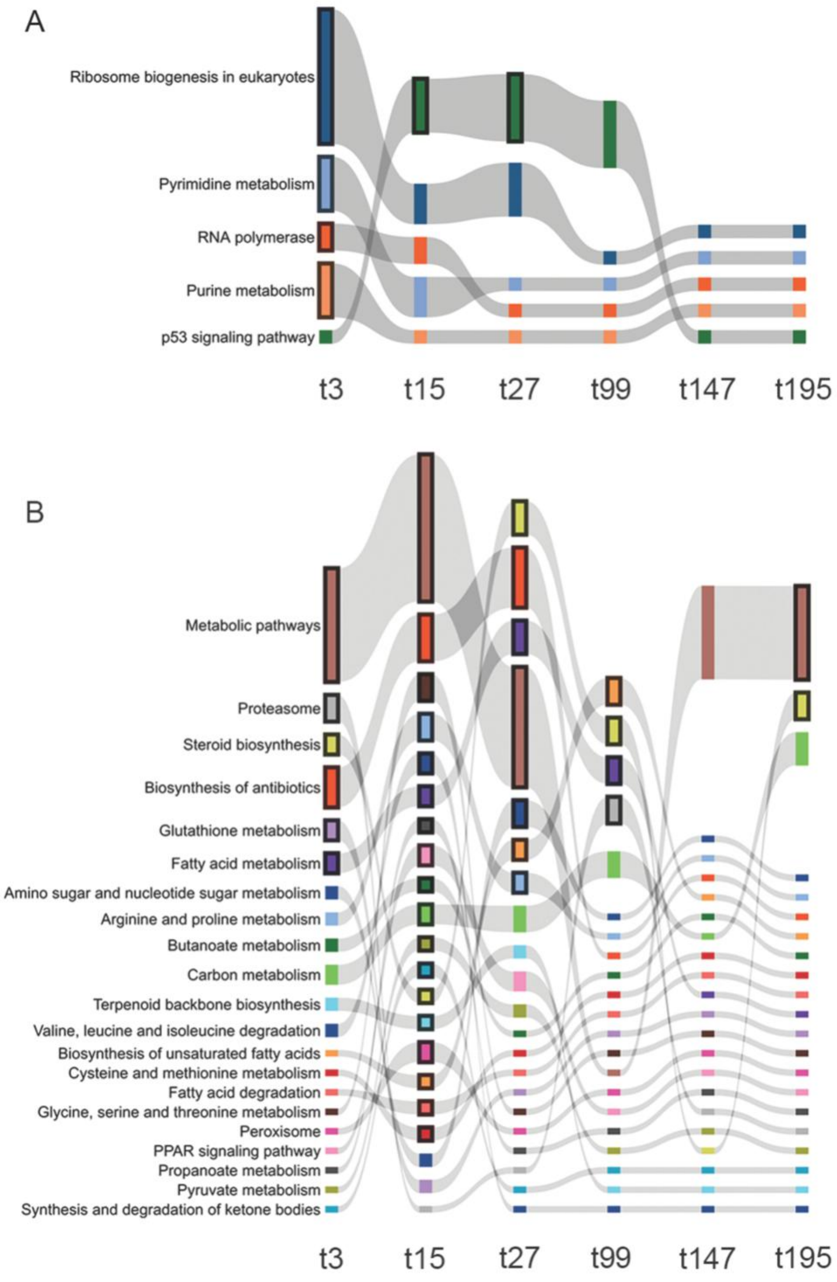


**Figure S2: Temporal map of up regulated and down regulated processes during liver regeneration.** The 1265 DEGs were categorised into up regulated (A) and down regulated (B), according to their expression levels in t3 compared to t0, for ease of visualization. The functional annotation of statistically overrepresented processes was performed on the two categories separately. Using Sankey software, the processes were plotted in an alluvial plot. The height of the node represents percentage of genes observed in that process. The nodes with solid borders represent the significance of the process at the respective time point (Benjamini corrected $p<0.05$).

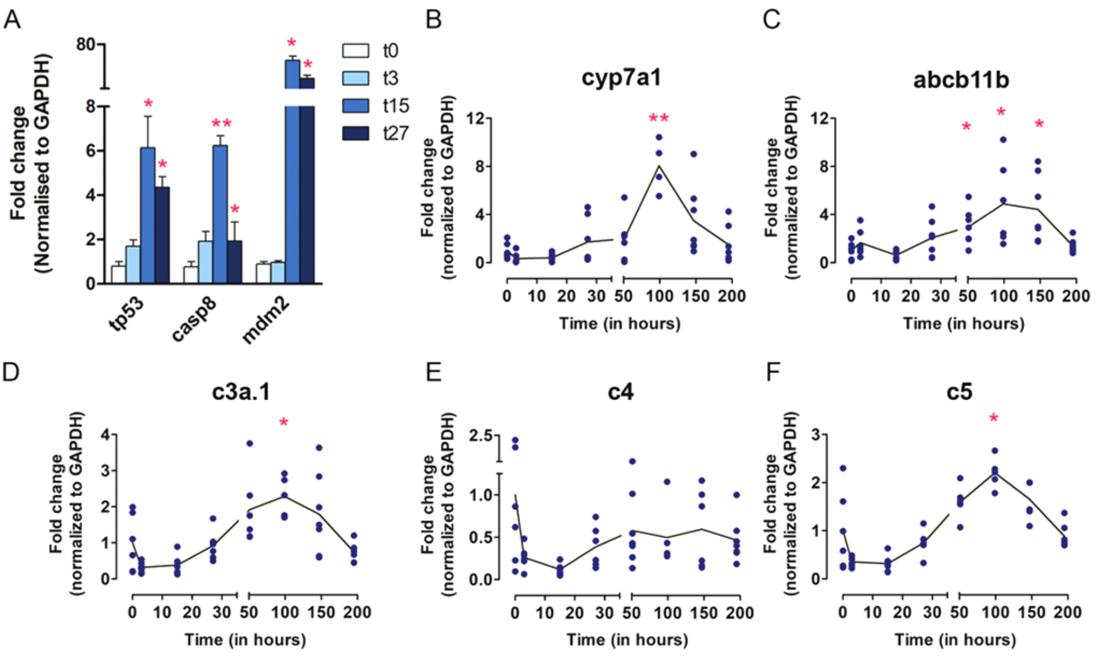


**Figure S3: Validation of DEGs in RNAseq by quantitative real time PCR.** Data was obtained from minimum of 4 and maximum of 7 independent adult zebrafish livers (biological replicates) for each time point. Ct values were normalized to GAPDH. (A) Gene expression changes for apoptosis genes, tp53, caspase 8 and mdm2. Fold changes by qRT PCR for (B) cyp7a1 (C) abcb11b (D) c3a.1, (E) c4 and (F) c5 were determined at all time points. Statistical significance was calculated using unpaired student t test (*p < 0.05, **p < 0.01) with respect to t0.

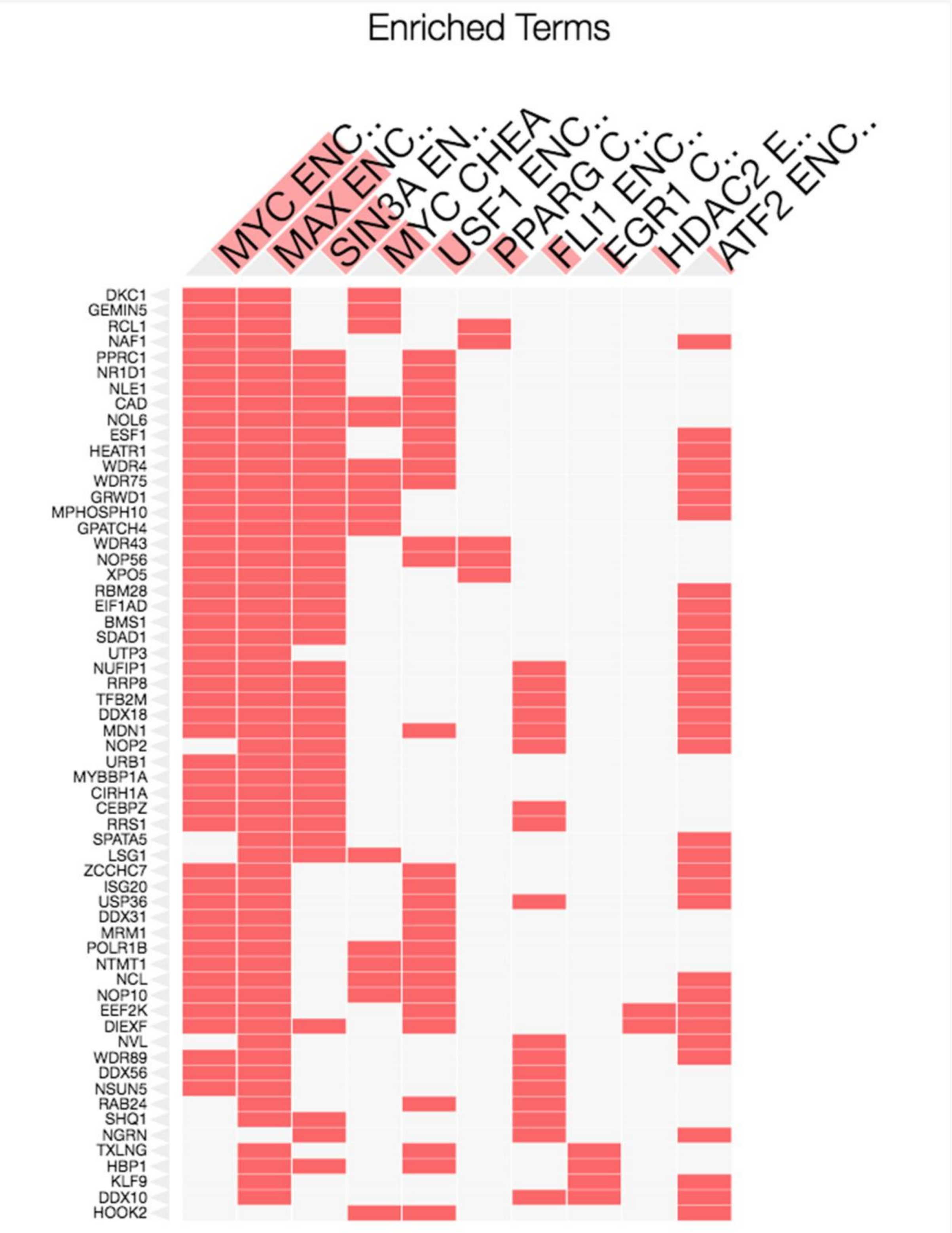


**Figure S4: ENRICHR analysis for transcription factors targets in the DEGs in module 2 and 4.** Combined gene list of modules 2 and 4 were analysed on the ENRICHR website. The X-axis shows the significantly enriched transcription factors for which target sites exist on the genes listed on the Y-axis. The TFs with lowest p value are highlighted with orange block.
